# Co-option of endogenous retroviruses through genetic escape from TRIM28 repression

**DOI:** 10.1101/2022.06.22.497016

**Authors:** Rocio Enriquez-Gasca, Poppy A. Gould, Hale Tunbak, Lucia Conde, Javier Herrero, Alexandra Chittka, Robert Gifford, Helen M. Rowe

**Author notes:** Equal Contribution (Rocio Enriquez-Gasca, Poppy A. Gould).

## Abstract

Endogenous retroviruses (ERVs) have rewired host gene networks through co-option of their enhancers. To explore which ERVs get co-opted and why, we tracked the epigenetic fate of murine IAPEz elements using an *in vitro* model of embryonic stem cell (ESC) to neural progenitor cell (NPC) differentiation. TRIM28-repression depended on a 190bp sequence, previously shown to confer IAPEzs with retrotransposition activity. A subset of IAPEzs (∼15%) exhibit genetic divergence from this sequence, which we term escapees. While repressed IAPEzs succumb to a previously undocumented epigenetic handover from H3K9me3 in ESCs to H3K27me3 in NPCs, escapee IAPEzs evade repression, resulting in their transcriptional derepression in NPCs. Escapee IAPEzs enhance expression of nearby neural genes and contribute to gene expression differences between mouse strains, which we discern by employing IAPEz insertion polymorphisms. In sum, co-opted ERVs stem from genetic escapees that have lost vital sequences required for both TRIM28 restriction and autonomous retrotransposition.

**HIGHLIGHTS:**

- Tracking the epigenetic fate of ERVs through neural differentiation reveals which ERVs are subject to co-option and why.
- Most ERVs succumb to H3K9me3 deposition in ESCs with H3K27me3 memory in NPCs.
- ERVs that act as enhancers to nearby neural genes have undergone genetic escape from TRIM28 repression.
- Epigenetic evasion is an evolutionary trade-off that comes with loss of sequences necessary for retrotransposition.

Graphical abstract.

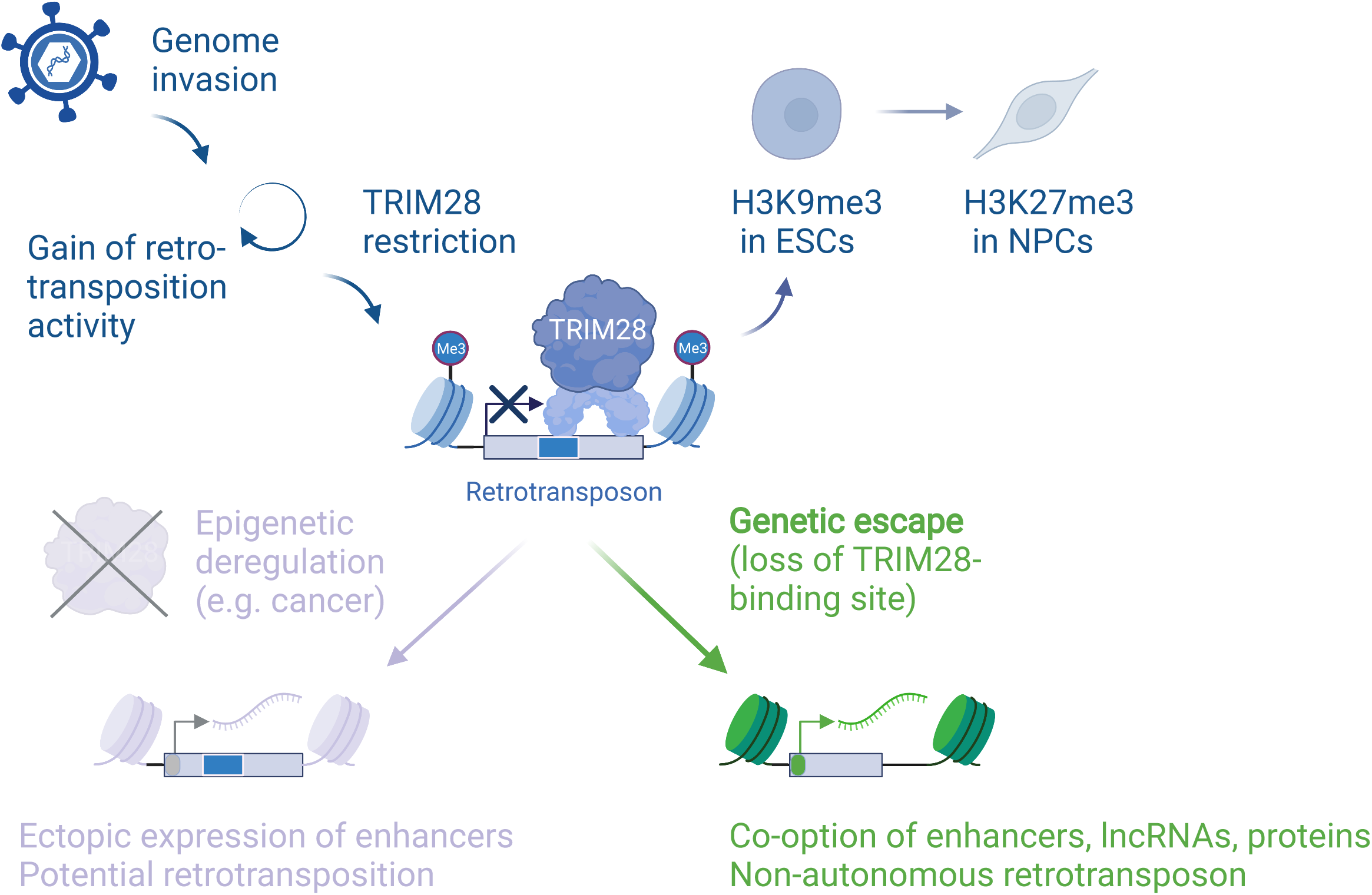

## INTRODUCTION

Mammalian genomes are constantly co-evolving with the abundant transposable element (TE)-derived DNA burden that they are comprised of (de Koning et al., 2011). TEs possess their own functional sequences, which can be repurposed by the host in a process known as co-option, for example to rewire gene regulatory networks (Chuong et al., 2016) or generate novel proteins (Cornelis et al., 2017; Mi et al., 2000). They also contain sequences that are targeted by TRIM28-mediated epigenetic silencing to protect genome integrity (Jacobs et al., 2014; Rowe et al., 2013a; Wolf et al., 2015). This duality represents an evolutionary dilemma in terms of how and when a TE becomes silenced vs. co-opted to affect a cellular function.

Several prominent examples of co-option of TE-derived sequences in different molecular roles have been described (Cosby et al., 2019; Enriquez-Gasca et al., 2020; Mi *et al*., 2000). Neural lineages represent a fertile ground for innovation with neocortex-specific enhancers conserved in present-day mice that are derived from ancient TEs dating back to amniote genomes (Notwell et al., 2015). Intriguingly, somatic mosaicism resulting from L1 activity has been documented in the human brain (Muotri et al., 2005; Sanchez-Luque et al., 2019) potentially contributing to phenotypic variability. A neuronal protein, *Arc* is derived from a retroviral *Gag* gene and functions as a viral-like capsid, transferring mRNAs from neuron to neuron (Pastuzyn et al., 2018). Despite well-documented cases of TE co-option, however, understanding the early events promoting this process remains a challenge. Here, we set out to pinpoint how co-option events may emerge by interrogating which endogenous retroviruses (ERVs) gain active enhancer activity and why. Enhancer activity serves as a proxy for subsequent co-option events including rewiring of host genes through cis-regulatory elements (Chuong *et al*., 2016), as well as the generation of novel long noncoding RNAs (Lu et al., 2014) and proteins derived from ERVs (Grow et al., 2015; Ng et al., 2019). We employ mouse embryonic stem cells (ESCs) as a developmental model and focus on an actively transposing murine ERV family, the Intracisternal A-type particles (IAPEz with long terminal repeats, of the LTR1/1a type)(Qin et al., 2010; Rebollo et al., 2020) to map evolutionarily-recent gain-of-enhancer events.

We map TRIM28-repression to overlap the signal peptide that targets IAPEz particles to the endoplasmic reticulum and which has been shown to have conferred IAPs with the ability to retrotranspose (Fehrmann et al., 2003; Magiorkinis et al., 2012; Ribet et al., 2008). Tracking the epigenetic fate of endogenous IAPEz elements reveals that those with the TRIM28 binding site are laden with H3K9me3 in ESCs and H3K27me3 in NPCs. A minority of IAPEz copies exhibit sequence divergence at the TRIM28 binding site, which mirrors their transcriptional derepression in NPCs, and we hereafter refer to these integrants as ‘escapees’. Escapee IAPs act as enhancers to nearby genes, which we discern by focusing on interstrain polymorphic IAPEz insertion sites. From this pool of new enhancers, only those with beneficial effects on adjacent genes would be targeted by natural selection, whereas the rest may reside as neutral events subject to further decay. Taken together this work shows that epigenetic silencing depends on TRIM28 targeting of vital sequences needed for retrotransposition. Genetic escape from epigenetic silencing paves the way towards ERV domestication. This in turn explains why the noted downregulation of epigenetic complexes in some cancers (Tunbak et al., 2020), can potentially unveil rare retroelement copies intact for retrotransposition (Rodriguez-Martin et al., 2020).

## RESULTS

### TRIM28 targets and represses IAPEz elements independent of the YY1 and PBS sites

We explored the early steps of co-option by focusing on IAPEz elements, which adapted to retrotranspose through gain of an endoplasmic reticulum (ER)-targeting signal at the N-terminal of GAG and loss of their retroviral envelope gene (Figure 1a – top panel). We define full-length IAPEz LTR1/1a integrants in the reference C57BL/6J genome (referred to hereafter as IAPEz) as those that (1) contained two LTRs of the relevant subtype (LTR1/1a), (2) possessed two LTRs in the same orientation, (3) were separated by less than 20kb and (4) were annotated to have ‘IAPEz’ internal sequences. We then cross-referenced these positions with the reported list of deletions in the 129P2/Ola genome to define a list of 838 full-length IAPEz copies known to be present in both C57BL/6J and 129 mouse strains (Figure 1a – bottom panel, Supplementary Information 1).

**Figure 1.**
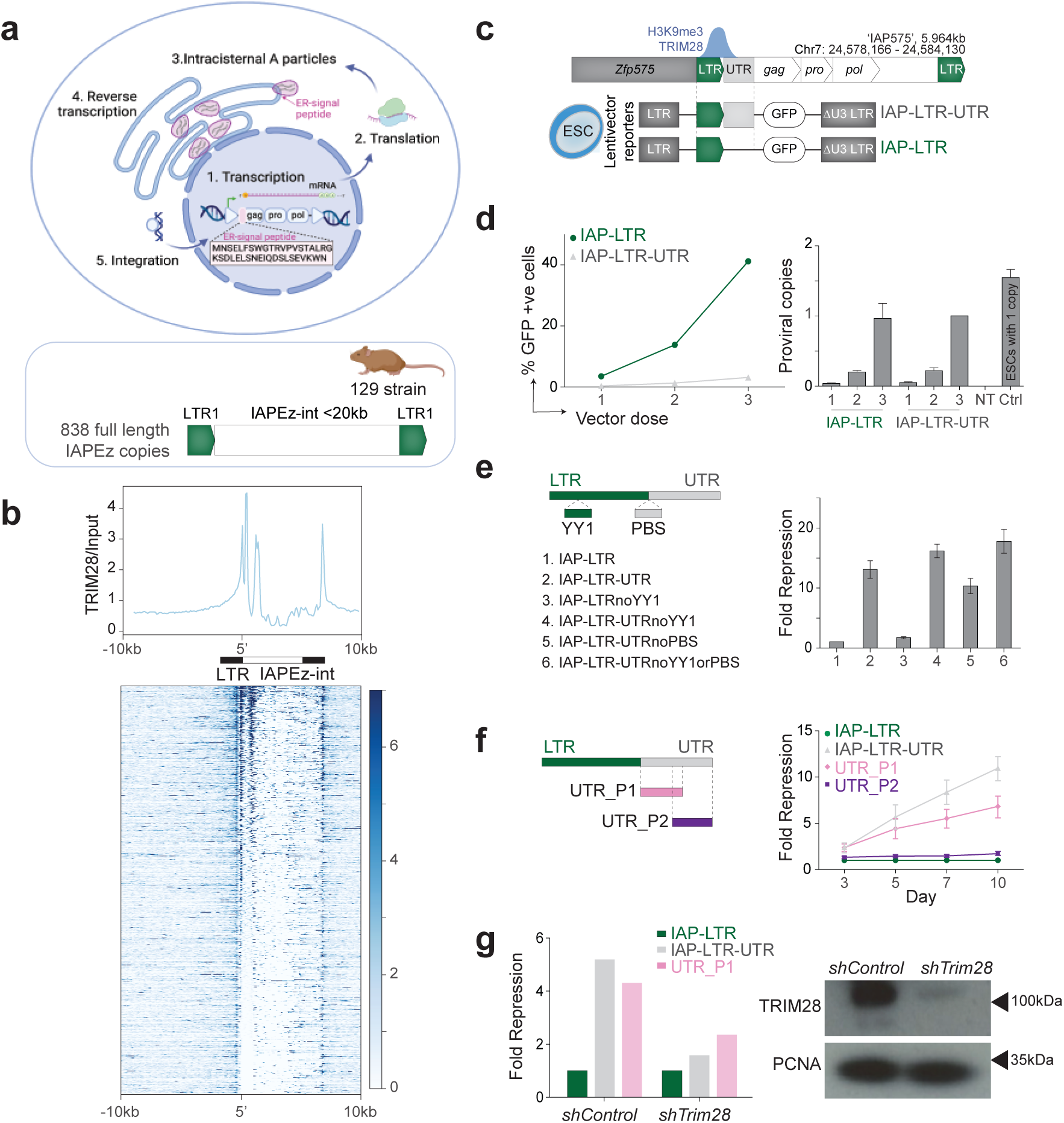
TRIM28 targets and represses IAPEz elements independent of the YY1 and PBS sites. (a) Retrotransposition cycle of IAPEz elements. IAP elements gained an endoplasmic reticulum (ER) signal peptide upstream and in frame with GAG, retargeting particles to the ER (top). Schematic representation of the sequence structure of IAPEz which was used to annotate full elements in this study (bottom). (b) TRIM28 fold enrichment to total input over full-length IAPLTR1/1a elements, where the 3’ end coordinate of their 5’ LTR was used as the reference point. Signal is represented as a profile plot (top) or a heatmap sorted by TRIM28-binding intensity (bottom). (c) Schematic representation of a TRIM28-repressed IAP in chromosome 7 (IAP575) (top) and of the lentivector reporter constructs with the LTR +/- its endogenous 5’UTR and GAG junction (bottom). (d) Percentage of GFP positive (+ve) cells by vector dose in ESCs transduced with the reporter construct in (c), results are shown for day 8 post transfection (left); relative number of proviral integrants for both constructs (right). (e) Fold repression of reporters normalised to expression of IAP-LTR in ESCs where constructs harbour a deletion of the YY1 binding site or the PBS or both (depicted left), data shown for day 8 post transduction (right). (f) Fold repression of the reporter normalised to expression of IAP-LTR in ESCs for constructs comprising part 1 (P1) or part 2 (P2) of the 5’UTR as shown left, in addition to constructs shown in (c), over a timecourse of three to ten days (right). (g) Fold repression of indicated constructs in control or *Trim28* depleted (shRNA) ESCs, data from day 3 post transduction before TRIM28 depletion is lethal (right). Western blots showing successful knockdown of TRIM28 with PCNA as a loading control.

We employed public ChIP-seq data from (De Iaco et al., 2017) to visualise the binding intensity of TRIM28 on these 838 IAPEz LTRs and 10kb flanking regions. One binding site for TRIM28 corresponded to the LTR and was present in full-length (Figure 1b) and solo LTRs (Supplementary Figure 1a), while a second prominent TRIM28 peak mapped to the IAPEz UTR of full-length elements (Figure 1b). Cloning sequences of an IAPEz integrant (termed IAP575, Figure 1c) located downstream of the gene *Zfp575,* at which we previously detected TRIM28, SETDB1 and repressive H3K9me3 to be enriched at (Rowe et al., 2013b), showed that the LTR-UTR and not the LTR alone was sufficient to confer repression in a reporter assay in ESCs (Supplementary Figure 1b, Figure 1d). Silencing initiated by the UTR was detected by day 3 post transduction (Supplementary Figure 1c) and complete by day 8. We verified that reporter repression was not due to lack of vector integration (Figure 1d) and 3T3 cells, which cannot establish heterochromatin served as an additional control. As expected, both constructs were equally expressed in these cells (Supplementary Figure 1d). This suggested the TRIM28 binding site within the IAPEz UTR to be a key determinant of epigenetic repression. The IAPEz LTR harbours a YY1 binding site like LINE-1 elements, which have recently been shown to depend on this site for their epigenetic repression (Sanchez-Luque *et al*., 2019), whereas the retrovirus MLV is silenced through its PBS (Wolf and Goff, 2007) and YY1 site (Schlesinger et al., 2013). We therefore asked if the YY1 site or PBS site were necessary for repression individually or in combination and found both to be dispensable (Figure 1e, Supplementary Figure 1e). Subsequent reporter assays mapped repression to the proximal part of the IAPEz UTR (Figure 1f, Supplementary Figure 1f), which was relieved upon TRIM28 *shRNA*-mediated depletion of TRIM28 (Figure 1g, Supplementary Figure 1g), as expected.

### Epigenetic repression maps to an 190bp sequence overlapping the IAPEz signal peptide

In order to more precisely map the sequence required for TRIM28-repression, we drew on previous sequence annotation of the IAPEz UTR. The IAPEz UTR contains two direct repeats (DRs) (Mietz et al., 1987) between which resides an ER targeting peptide that is in frame with GAG and which has been shown to have conferred an infectious IAPEz progenitor with the *de novo* ability to retrotranspose (Ribet *et al*., 2008). The remnants of DRs suggests that this signal peptide, which derives from host sequences could have been captured itself through a retrotransposition event. Deletion of DR1 (96bp) or DR1 plus a fraction of DR2 (127bp) showed that the 127bp deletion was sufficient to relieve reporter repression, indicating that DR1 is necessary for repression (Figure 2a, Supplementary Figure 2a). To establish whether DR1 is also sufficient to confer repression or whether DR1 plus DR2 is required, we tested the repressive effect of these sequences upstream of the LTR reporter construct, in either sense (S) or antisense (αS) orientation. This revealed that DR1 and DR2 in combination are necessary and sufficient to establish repression to the same degree as the whole UTR sequence (Figure 2b, Supplementary Figure 2b). Thus, we have defined a novel 190bp sequence encompassing both DR1 and DR2 as the functional repressor of IAPEz elements.

**Figure 2.**
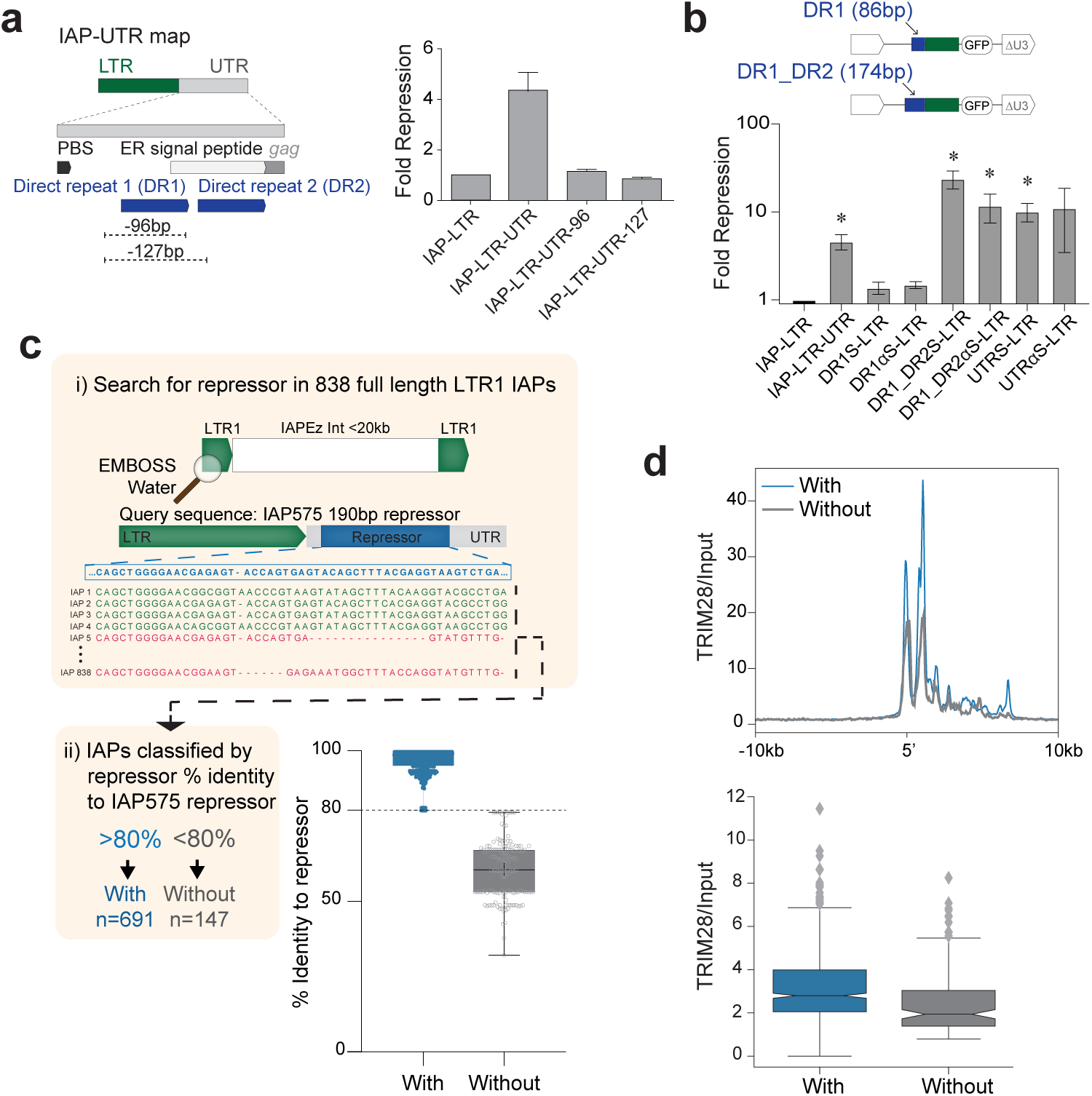
Epigenetic repression maps to an 190bp sequence overlapping the IAPEz signal peptide. (a) Schematic representation of the first half of the 5’ UTR showing the direct repeats and host-derived ER-targeting signal peptide together with the 96 or 127bp deletions (left) assayed in ESCs for fold repression as previously (right). (b) Fold repression in ESCs of reporter constructs containing one or both direct repeats upstream of the IAP LTR promoter in either orientation (significance determined with two-tailed paired t tests, IAP-LTR-UTR p=0.0162; DR1_DR2S-LTR p=0.0184; DR1_DR2aS-LTR p=0.0496; UTRS-LTR p=0.0226). (c) Depiction of the strategy used to classify sequences of full-length IAPEz elements by their percentage identity to the 190bp functional repressor sequence of IAP575. Boxplots represent 1st and 3rd quartile, where the central line corresponds to the median; whiskers are ×1.5 of the interquartile range. (d) TRIM28 signal normalised to total input for IAPEz elements separated by their identity to the functional repressor shown as a profile plot (top) or boxplot where the mean signal across the 20kb interval depicted above is taken for each element (bottom). Mann-Whitney U test p-value=1.7e-11.

Having identified the functional repressor sequence, we determined its conservation across the 838 full-length copies of IAPEz present in the genome of 129P2/Ola mice. We classified IAPEz copies based on their percent identity to the canonical IAP575 repressor sequence, with those exhibiting 80% or greater identity classified as an IAPEz ‘with’ a functional repressor and those with <80% sequence identity as an IAPEz ‘without’ a repressor, reasoning that these two groups may exhibit differences in their epigenetic fate (Figure 2c). 82% of IAPEz elements contained a full repressor, which we term canonical IAPs, while only 18% fell into the without group, in which there was a wide range of percent-divergence from the repressor between individual integrants (Figure 2c). To address whether genetic sequence divergence from the repressor has functional consequences, we compared the TRIM28 binding between the with- and without-repressor IAP groups. This analysis unveiled a much stronger TRIM28 signal for the canonical IAPEz copies (Figure 2d) indicating that sequence divergence may alter the epigenetic regulation of these elements.

### The epigenetic fate of IAPEz ERVs in neural progenitor cells is determined by their sequence

To track the epigenetic consequences of the genetic changes we observed at the repressor sequence, we utilised an *in-vitro* system of ESC to neural progenitor cell (NPC) differentiation and CUT&RUN chromatin profiling (Figure 3a). While IAPEz elements are largely identical to one another, preventing the detection of uniquely-mapping short reads from CUT&RUN sequencing data within them, H3K9me3 enrichment at these elements spreads into flanking regions (Rebollo et al., 2011). This phenomenon allows these elements to be mapped at the subfamily level using either multi-mapping (Figure 3b) or uniquely-mapping reads (Supplementary Figure 3a). H3K9me3 enrichment was associated with integrants containing a functional repressor sequence, consistent with the pattern of TRIM28 binding observed in ESCs. This is in line with the described TRIM28 regulation of ERVs in NPCs (Fasching et al., 2015). Interestingly, H3K9me3 spreading was prominent in ESCs but not in NPCs with repressor-less IAPEzs exhibiting the lowest levels of H3K9me3 (Figure 3b). This suggests that IAPEzs can invoke repression on nearby genes in early development, whereas in NPCs there is less spreading of repression and a more favourable context for epigenetic escape of the repressor-less IAPEz copies (Figure 3b, middle panel).

**Figure 3.**
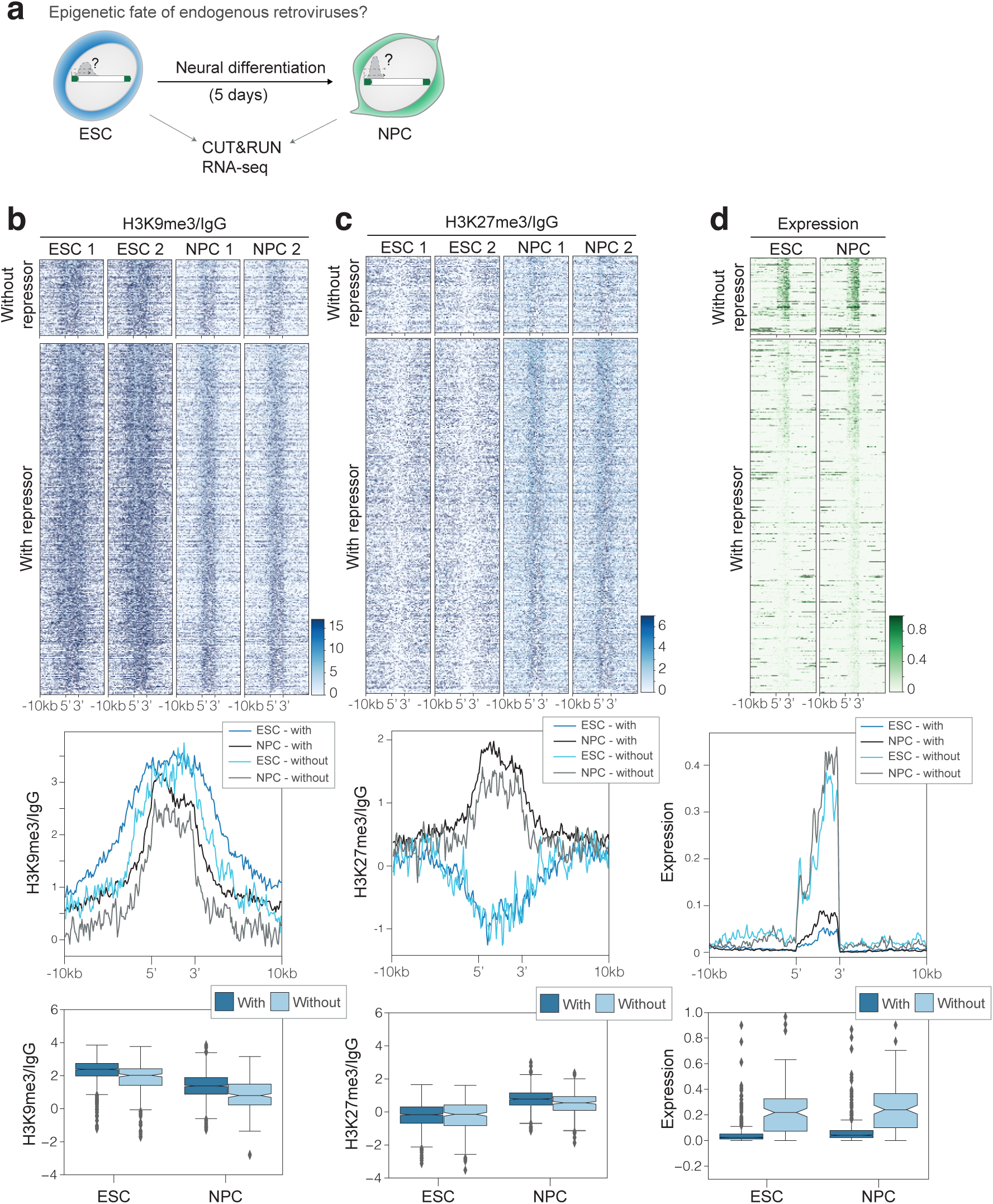
The epigenetic fate of IAPEz ERVs in neural progenitor cells is determined by their sequence. (a) Schematic of ESC to NPC differentiation. (b) H3K9me3 CUT&RUN signal normalised to IgG across 838 IAPEz elements and surrounding 10Kb sequences to either end, separated by the presence of the repressor (with and without) and sorted by decreasing levels of intensity as a heatmap for two replicates (top). Profile plots depict the trimmed mean across all elements in the indicated category, across 100bp bins relative to the start and end coordinates of the element, where each full IAP is depicted in 50 bins (middle) and boxplots show mean signal across the element and flanking regions for each IAPEz. Mann-Whitney U test, FDR corrected P-value for ESC p=2.4e-19; NPC p=6.4e-21. (c) H3K27me3 CUT&RUN signal normalised to IgG for two replicates as in (b), where elements in the heatmap are sorted by the decreasing H3K9me3 signal. Mann-Whitney U test, FDR corrected P-value value for ESC p=0.94; NPC p=1.6e-09. (d) Mean RNA-Seq signal across 3 replicates and normalised to number of mapped reads. Rows in the heatmap are sorted by decreasing H3K9me3 signal. Boxplots show mean signal across the element and 1000bp flanking regions for each IAPEz. Mann-Whitney U test, FDR corrected p value for ESC p=4.6e-08; NPC p=7.8e-08.

The significant loss of H3K9me3 at repressor-less IAPEzs in NPCs prompted us to measure the levels of H3K27me3, which is deposited by the PRC2 complex and associated with repression of cell type-specific genes (Mohn et al., 2008). Strikingly, we saw significant levels of H3K27me3 at IAPEz elements in NPCs, which was completely absent in ESCs (Figure 3c, Supplementary Figure 3b). While H3K27me3 has been documented to safeguard IAP silencing in germ cells (Huang et al., 2021) and upon induced loss of DNA methylation in ESCs (Walter et al., 2016), an epigenetic switch from H3K9me3 to H3K27me3 at IAPs has not been reported during NPC differentiation and further highlights IAPEzs as dynamically regulated sequences in neural development. We next looked at RNA expression of IAPEz elements in both cell types, as a proxy for the potential emergence of co-option events. Results showed unequivocally that a decrease in silent epigenetic marks at escapee IAPEzs correlated with their transcriptional derepression (Figure 3d, Supplementary Figure 3c). Escapee IAPEzs exhibited a higher level of transcriptional activity in NPCs than ESCs, suggesting that co-option might be favoured in neural lineages.

### The IAP LTR1 U3 region is a potent enhancer and activates nearby neural genes

The transcriptional activation of escapee IAPEz copies posed the exciting possibility that some of these integrants may harbour intact enhancers that could activate host genes. Indeed, as IAPs are derived from retroviruses, they have a putative enhancer in the U3 region (Mietz *et al*., 1987; Zierler et al., 1992). We asked if the IAP U3 could act as a classical enhancer by cloning the IAP575 U3 sequence into a reporter construct upstream of a minimal SV40 promoter in sense and antisense configurations. Results illustrate that the U3 sequence, and the minimal putative 47bp enhancer sequence within it, function as potent enhancers in a cell-type independent manner (Figure 4a, Supplementary Figure 4a).

**Figure 4.**
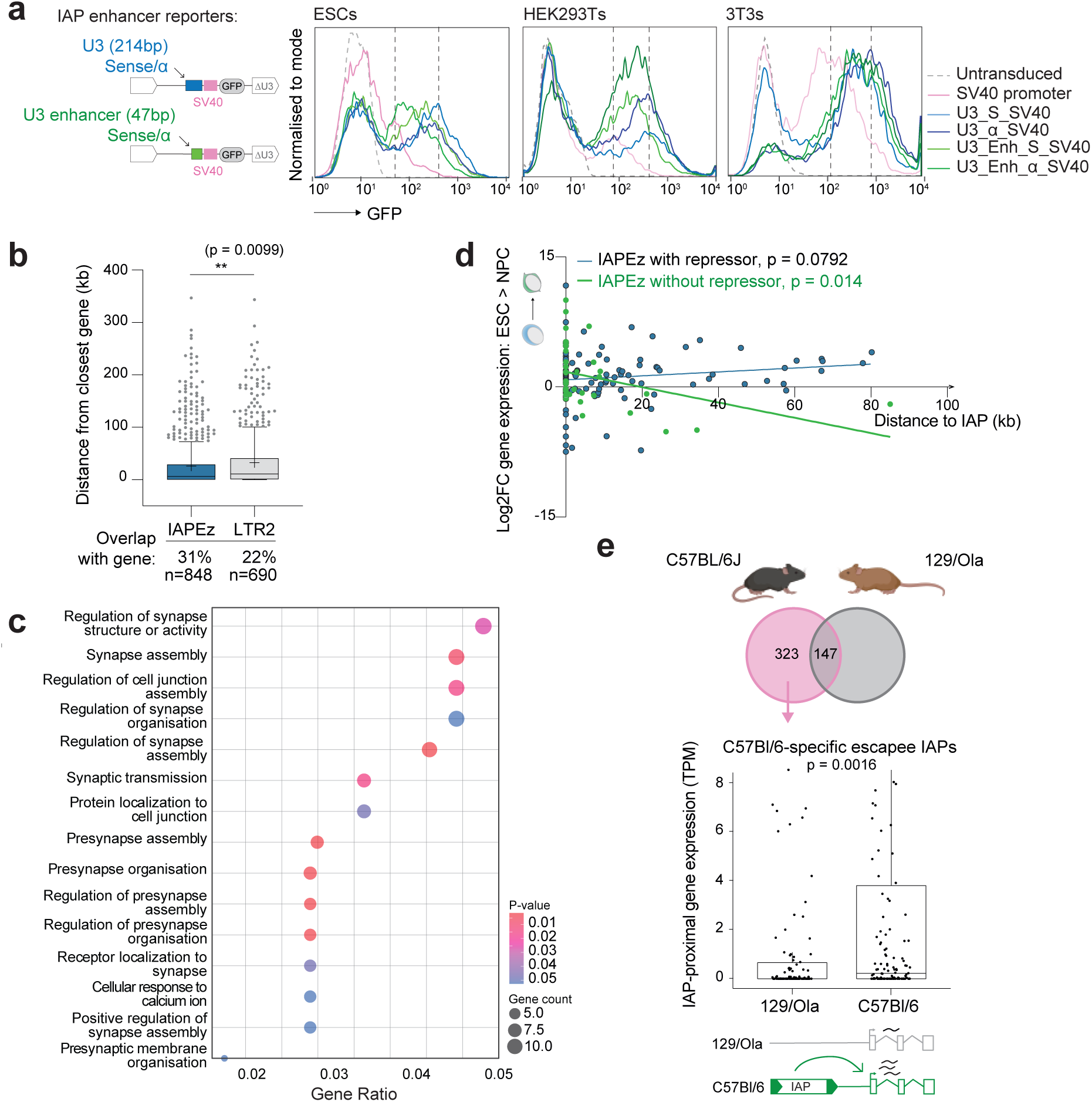
The IAP LTR1 U3 region is a potent enhancer activating nearby neural genes and accounting for interstrain gene expression differences. (a) GFP mean fluorescent intensity (MFI) of the stated cell lines transduced with reporter constructs (shown left) containing either the entire IAP U3 region or the minimal enhancer (47bp) within the U3, in sense and antisense orientations, upstream of a minimal SV40 promoter reporter that lacked its own enhancer. (b) Boxplot of distances between younger (IAPEz) and older (IAPLTR2/2a/2a2/2b) IAPs and their closest gene in kb, P-values calculated using Welch’s t-test; percentage of elements overlapping a gene as well as total number in each class is shown below plot. (c) Gene-set enrichment analysis to Gene Ontology annotations for genes closest to IAPEz elements. (d) Scatterplot depicting distance to closest gene and log2FoldChange in expression between NPCs and ESCs for IAPEz, subdivided into those with and without repressor. Linear regression and P-values shown. (e) Boxplot depicting TPMs of 323 genes closest to IAPEzs lacking a repressor sequence (escapee IAPs) which are present only in the C57Bl/6 strain, where expression values are for 129/Ola NPCs or C57Bl/6 Cortical Neurons from (Bonev *et al*., 2017). P-values were calculated using a paired Wilcoxon test.

With the aim to investigate whether escapee IAPEzs can act as enhancers for nearby genes, we first observed that a significant fraction (31%) of IAPEz elements overlap a gene, compared to 22% of IAPs with an IAPLTR2 type LTR, which we employed as an older IAP family for comparison. In terms of distance, IAPEz elements were also significantly closer to genes than their older IAPLTR2 counterparts (Figure 4b), further implicating young IAPEz elements as candidates for co-opted gene regulatory functions. When classifying the IAPEz-proximal genes by gene type, we found their relative proportions to be largely comparable to the whole genome, except for a slight but significant depletion for protein coding genes (Supplementary Figure 4b). However, when looking at the function of IAPEz-proximal genes, we uncovered them to be enriched in synapse-associated terms (Figure 4c), unlike the older IAPLTR2-proximal genes, which showed no enrichment of functional terms.

Considering both the enrichment of neural-related terms in IAPEz-proximal genes concurrent with the loss of heterochromatin at escapee IAPs in NPCs, we asked whether there was an association between changes in gene expression upon NPC differentiation of ESCs (log2foldChange) and the distance to the closest IAPEz, depending on the presence or absence of the repressor sequence. This analysis revealed a statistically significant positive correlation between the log2foldChange (NPC/ESCs) and the distance to the closest repressor-less IAPEz. Genes proximal to canonical IAPs with the repressor showed the opposite trend, in contrast, potentially pointing to a dampening effect on the expression of nearby genes (Figure 4d). Intriguingly, escapee IAPEz copies are also significantly closer to and more frequently overlap genes than those with a repressor (Supplementary Figure 4c).

### The enhancer effect of escapee IAPEz’s is revealed by interrogating strain-specific insertions

Taking advantage of the natural variation of IAPEz insertion sites across laboratory mouse strains (Lilue et al., 2018), akin to a resource of natural genome-editing experiments, we asked whether these polymorphisms could lead to strain-specific gene expression differences. Using a list of 323 C57BL/6J-specific escapee full-length IAPEz elements (Figure 1a) compared to the 147 C57BL/6J-129P2/Ola-shared escapee IAPEz elements, we scored the expression levels of IAPEz-proximal genes in ESC-derived cortical neurons, using data from (Bonev et al., 2017). Escapee IAPEz copies in C57BL/6J were associated with significantly higher expression of proximal genes in this strain (Figure 4e), while genes proximal to shared escapees showed no significant differential expression (Supplementary Figure 4d). Taken together these data indicate that escapee IAPEz integrants exert an enhancing effect on proximal host gene expression.

### Significant genetic divergence of the IAPEz signal peptide underpins gain-of-enhancer events

To interrogate the genetic changes permitting escape from repression, we performed multiple sequence alignments of IAPEzs with and without the repressor and the IAP575 sequence; this analysis revealed that the main genetic divergence in the escapee sequences occurs in the ER targeting peptide region. Performing alignments to the amino acid sequence of the ER targeting peptide, revealed a striking difference in % identity separating the escapee and repressed IAPEzs (Figure 5a, Supplementary Figure 5a). A closer inspection of the multiple sequence alignments of the sequence corresponding to the repressor for all 838 IAPEz elements revealed two subcategories of “with repressor” IAPEzs. These were defined by whether the IAPEz, like IAP575, contained either a single copy (IAPEz with 1) or a duplication (IAPEz with 2) of a 33bp segment of direct repeat 2. Interestingly, the escapee IAPEzs could also be subclassified according to iterations of this segment – into those either containing a duplication (IAPEz without 1), a single copy (IAPEz without 2) or those lacking it entirely (IAPEz without 3) (Supplementary Figure 5b). Intriguingly, the subcategory with the most severe deletions with respect to the TRIM28 binding site (IAPEz without 3) exhibited the most marked mRNA expression (Figure 5b). This further supports our model that a gain in ERV transcriptional activity stems from genetic escape from epigenetic silencing. The early steps of co-option, therefore, involve fixation of an endogenous retroviral family in the genome followed by its gain in retrotransposition activity, the molecular basis of which is targeted by TRIM28. Subsequent genetic escape in the repressor sequence enables the ERV regulatory sequences to gain activity while losing/decreasing their retrotransposition ability. Leading on from this, co-option of ERV enhancers, lncRNAs and novel chimeric proteins can emerge (Figure 5c).

**Figure 5.**
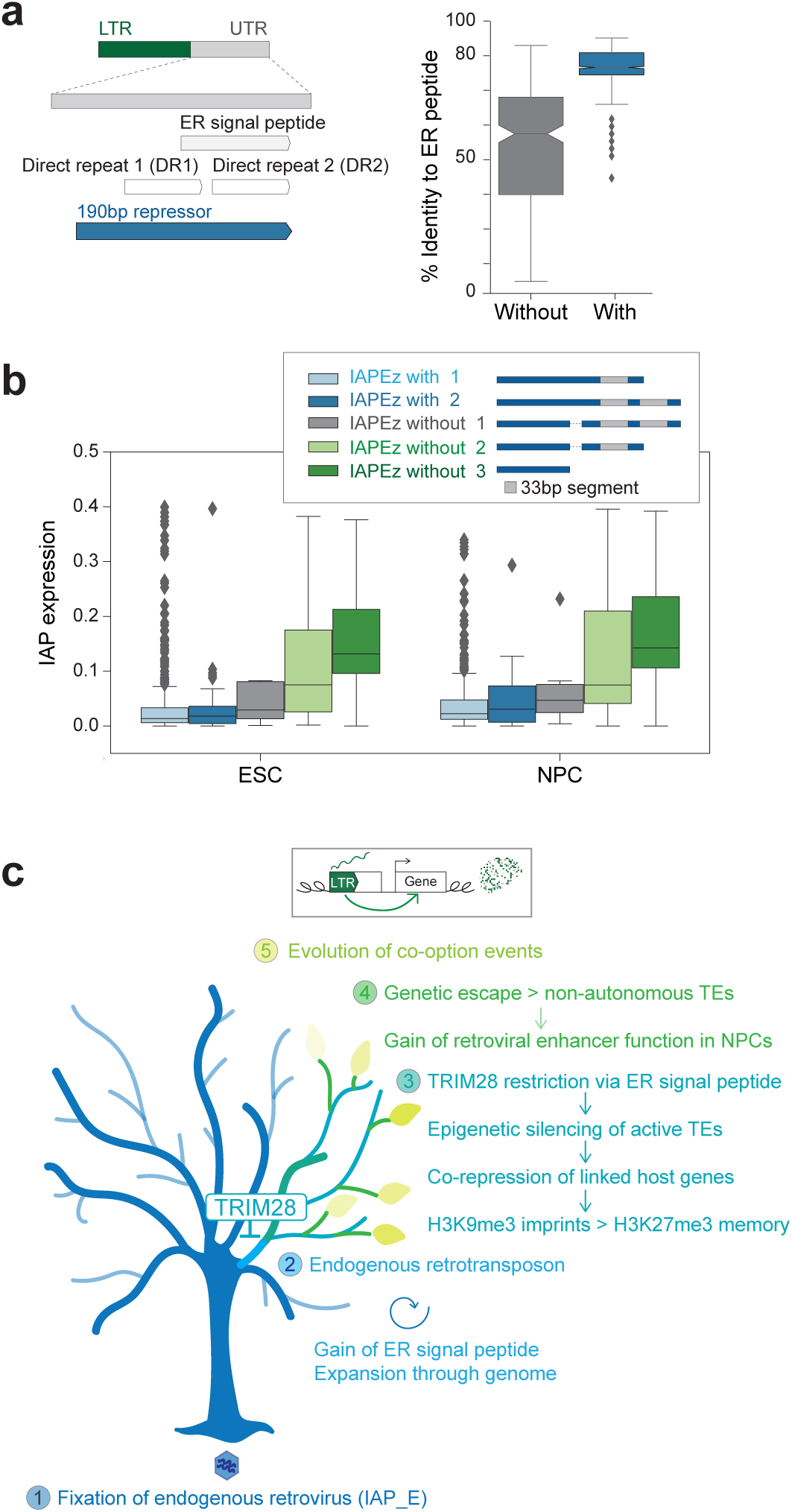
Dissection of genetic escape from TRIM28 repression and summary model of the proposed pathway to co-option. (a) Schematic of IAPEz UTR sequence depicting location of ER targeting peptide and two direct repeats relative to 190bp repressor (left). Boxplots of the percentage identity to the amino acid sequence of the ER targeting signal across IAPEz elements separated into with and without the repressor (right). (b) Boxplots of expression normalised to number of mapped reads, where IAPEz sequences have been further subdivided based on the observed changes in their repressor sequence (depicted in inset box). (c) Model. 1. An infectious retrovirus invades the germline and becomes fixed. 2. Evolution of an efficient retrotransposon through gain of a GAG ER-targeting signal derived from the host and loss of the envelope gene (Ribet *et al*., 2008). 3. Emergence of sequence-specific KZFP and TRIM28 restriction in early development for epigenetic repression of active retrotransposons through recognition of the ER-targeting signal. This involves co-repression of nearby genes through heterochromatin spreading of H3K9me3, which is consolidated with H3K27me3 memory following NPC differentiation. 4. Retrotransposon integrants that gain enhancer activity have undergone genetic escape through loss of the TRIM28 binding site, rendering them inactive TEs. Importantly, these retroviral enhancers are not active in mESCs, only following differentiation into NPCs. 5. Natural selection can then operate on ‘escapee’ retroviral enhancers and co-opt them for host gene expression. Escapee IAP retrotransposons represent a snapshot of evolution in action, since co-option events are ongoing in laboratory mouse strains. Gain of enhancer function in NPCs suggests that the brain represents a hotbed for retroviral co-option.

## DISCUSSION

In this work, we set out to identify the early steps paving the way to co-option of retrotransposons by focusing on IAPEz elements with an LTR1/1a as young and still actively retrotransposing ERVs in the mouse genome. In doing so, our study has highlighted the sequence corresponding to the ER targeting signal as a focal point of conflict between ERVs and their hosts. On the one hand, through previous genetic and biochemical investigation (Ribet *et al*., 2008), this sequence has been shown to have been vital to the endogenization of these elements, while on the other hand, here we identify it to represent a genetic vulnerability targeted by TRIM28 in the ongoing evolutionary arms race between TEs and their hosts. These results shed light on why epigenetic perturbations that cause reactivation of repressed retroelements, for example in cancer can not only lead to ectopic expression of ERV enhancers but also potentially to *de novo* retrotransposition (Gu et al., 2021). In the human genome, this would apply to LINE-1 elements, which are the only retroelements intact for autonomous retrotransposition (Rodriguez-Martin *et al*., 2020).

We uncover a subset of IAPEz elements exhibiting sequence divergence at the ER targeting signal, illustrating a selective pressure to escape from repression. We were able to identify at least three different versions of diverged sequences, suggesting that this escape has occurred more than once. By tracking the epigenetic state of IAPEz integrants through ESC differentiation to NPCs, we could demonstrate a functional effect of genetic escape: While canonical IAPs succumb to a dynamic regulatory pattern involving an initial silencing in ESCs by H3K9me3, with a handover to H3K27me3 in NPCs, repressor-less IAPEzs escape this regulation. We envision that canonical IAPEz elements are likely to be recognised by numerous KRAB-zinc finger proteins (KZFPs) targeting the UTR to initiate epigenetic repression via TRIM28, in a redundant manner (Imbeault et al., 2017; Wolf et al., 2020). Epigenetic derepression and the associated striking transcriptional activation are hallmarks of genetic escape that we identify as modifiers of the expression of nearby genes. We postulate that these escapees will promote future co-option events over long evolutionary timescales.

As we have focused on the early stages of co-option, most gain-of-enhancer events that we have documented here may be expected to be evolutionarily neutral or, indeed, deleterious. Expression of ERVs even if they are not full-length can lead to collateral damage. For example, ERV regulatory elements can unduly affect gene expression (Morgan et al., 1999) and cell fate (Macfarlan et al., 2012), and retroelement-derived nucleic acids can mimic viral replication intermediates and drive interferon responses and inflammation (Ahmad et al., 2018; Tunbak *et al*., 2020). Importantly though these changes are expected to be selected against and therefore not perpetuated, while beneficial acquired activity of ERVs will be co-opted and retained. Polymorphic IAPEzs have been documented to reside as metastable epialleles with variable DNA methylation levels (Kazachenka et al., 2018) and it is likely that the few integrants on their way to being co-opted to regulate host genes are those that are able to completely resist DNA methylation.

The proximity of IAPEz elements to genes enriched in neural functions is curious. Reasons for this could relate to the longer than average length of neural-related genes (Gabel et al., 2015; Sibley et al., 2015) or because this lineage is permissive to some degree of perturbation (Linker et al., 2017). In addition, and perhaps as a cause or consequence of the previous points, we have seen a gain of H3K27me3 at IAPEzs specifically in NPCs. The enrichment of this histone modification, which has been proposed to function as a placeholder of sequences to be activated later in neural development (Mohn *et al*., 2008), may point to another layer of gene regulation associated with IAPEz elements in NPCs. Although the strain-specific effects we have found suggest that some of these enhancer effects already have a functional impact on the expression of host genes, the history of IAP co-option in the mouse genome is still being written. Still, IAPEz elements appear to be remarkably well poised, both in terms of their genomic context and their epigenetic regulation, for co-option in neural tissues. Looking back at ancient co-option events that have been preserved throughout millions of years indeed reveals that ERVs have been notably coerced into co-option in neural lineages (Pastuzyn *et al*., 2018).

The emerging picture from our studies, here of an active retrotransposon highlights the ongoing battle between selective forces driving both the expression and further transposition of these sequences, and the host genome’s struggle to keep these genomic invaders in check. We envision that in the face of this evolutionary arms race, an ultimate compromise is likely to be struck, where these invading sequences can be repurposed for the benefit of the host and thus earn the genomic space that they have colonized.

## ACKNOWLEDGEMENTS

We thank Austin Smith for the Sox1-GFP reporter mESCs and James Briscoe for advice on NPC differentiation. We thank Connor Husovsky for technical help. We thank Miguel Branco and Pradeepa Madapura for helpful comments on the manuscript. We thank labs within QMUL Epigenetics Hub for reagents and advice and labs in the Blizard Centre for Immunobiology for advice. We thank members of the Rowe Lab, Liane P. Fernandes and James Holt, for helpful discussions. This work was funded through an ERC starting grant (678350, TransposonsReprogram) to HMR, supporting REG and PAG and a Sir Henry Dale Fellowship through the Wellcome Trust and Royal Society (Grant number 101200/Z/13/Z) awarded to HMR, which supported HT. HMR is funded by a Barts Charity Lectureship (MMBG1R).

## CONTRIBUTIONS

Experiments were performed by PAG, HT, AC and HMR. Data analysis was by REG, PAG, LC, JH, RG and HMR. Ideas were contributed by all authors. REG, PAG and HMR conceived the project and wrote the paper. All authors read and approved the final manuscript.

## INCLUSION AND DIVERSITY

One or more of the authors of this paper self-identifies as an underrepresented ethnic minority in science. One or more of the authors of this paper self-identifies as a member of the LGBTQ+ community.

## SUPPLEMENTARY FIGURE LEGENDS

**Supplementary Figure 1.**
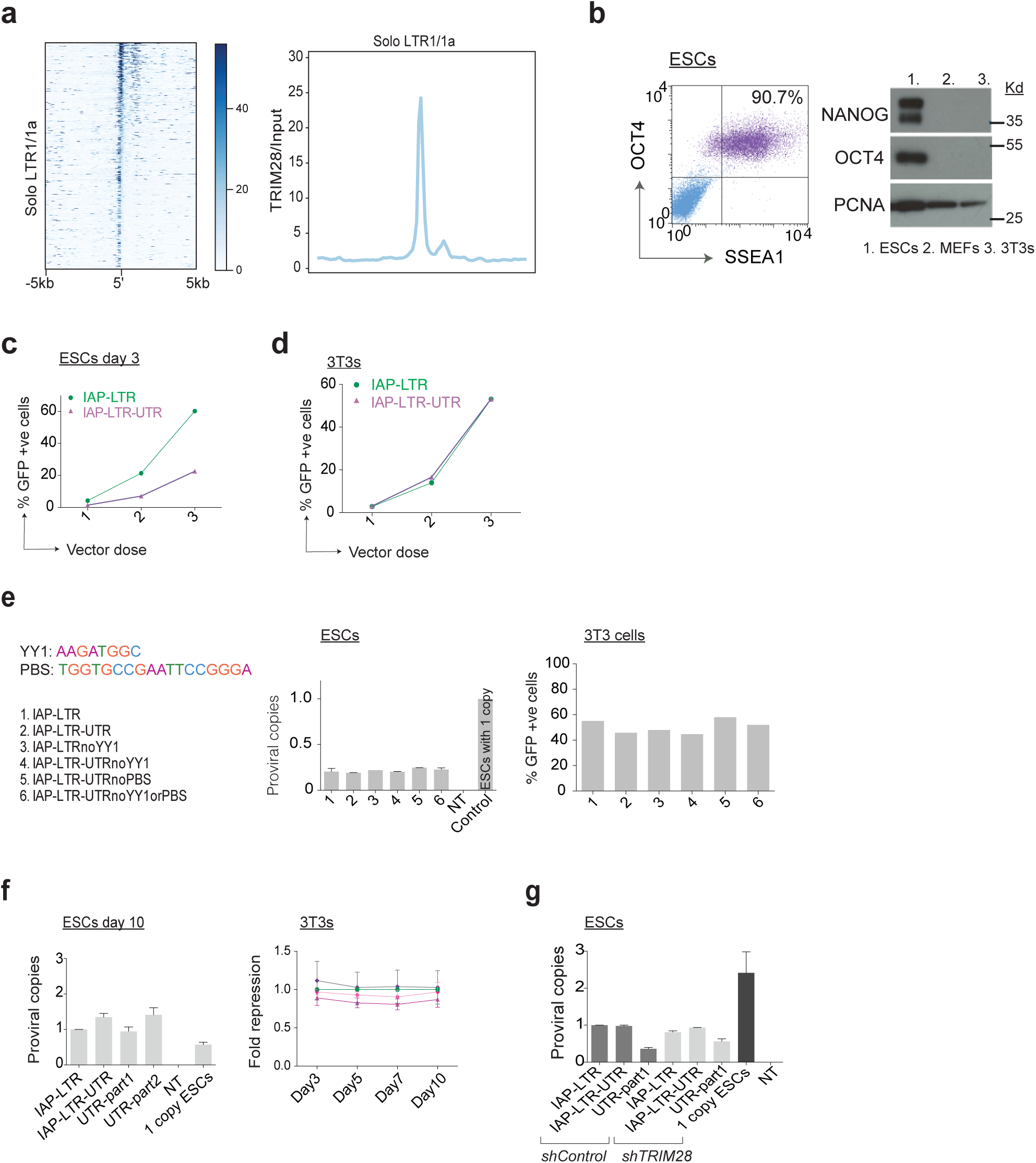
(a) TRIM28 fold enrichment relative to total input over solo-LTR IAPLTR1/1a elements, where the 3’ end most coordinate of their 5’ LTR was used as the reference point. Signal is represented as a heatmap sorted by TRIM28-binding intensity (left) or as a profile plot (right). (b) Mouse embryonic stem cells (ESCs) used for repression assays were verified to stain for OCT4 and SSEA1 by flow cytometry and for NANOG and OCT4 by Western blot. MEFs: mouse embryonic fibroblasts; 3T3s: NIH 3T3 immortalized fibroblasts. (c) Day 3 post transduction results for day 3 of reporter repression assay in ESCs shown for day 8 in Figure 1d. (d) Parallel reporter assay experiment to Figure 1d carried out in 3T3 cells. (e) Complementary data for Figure 1e to verify that all IAP reporter constructs tested here are expressed to similar levels in 3T3 cells as a control cell line and the YY1 and PBS sequences present in IAP575 are given (left). (f) Proviral copies of each construct used in Figure 1f in ESCs (left) and complementary data in 3T3 cells (right). (g) Proviral copies for each construct used in the *shControl* and *shTRIM28* ESC lines in Figure 1g.

**Supplementary Figure 2.**
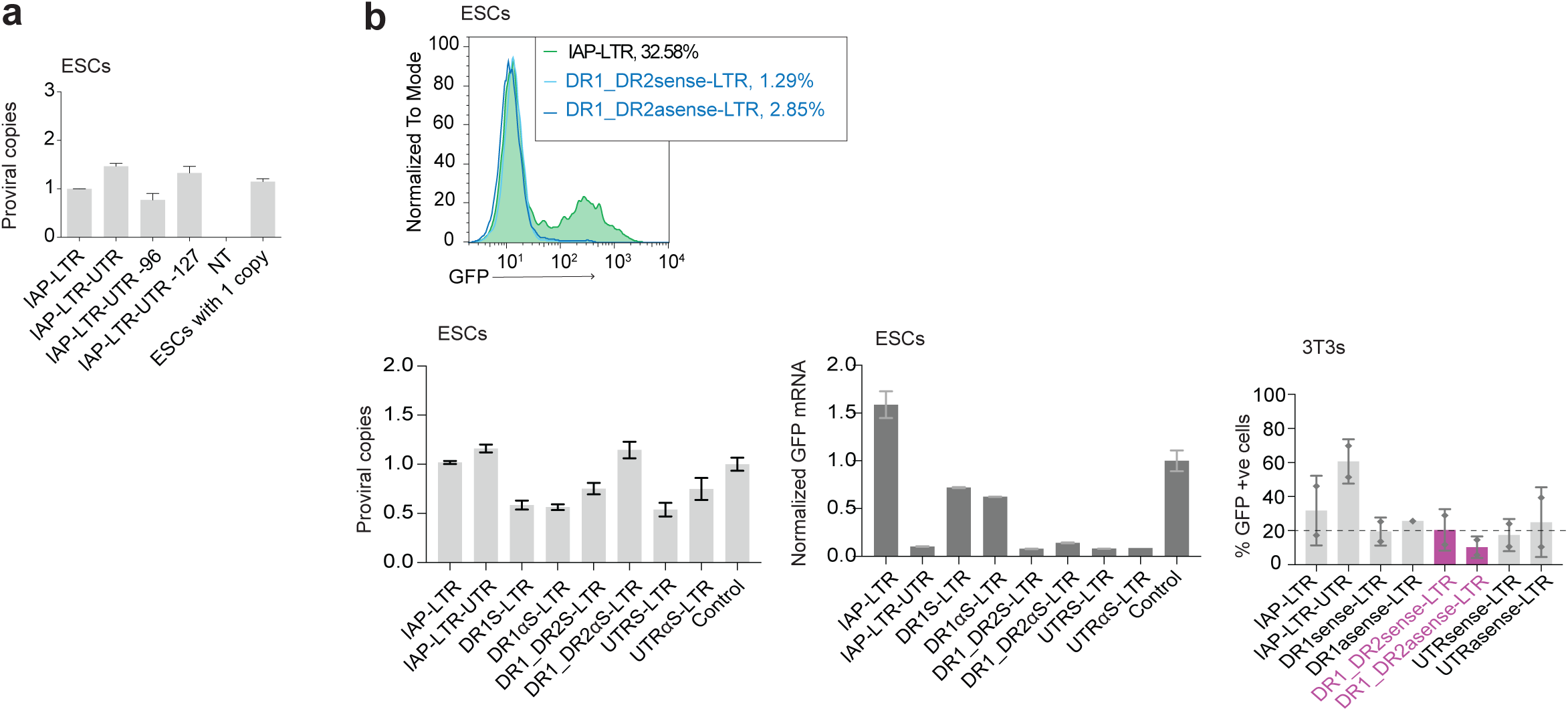
(a) Complementary integration data for Figure 2a verifying that epigenetic repression of IAP constructs was not due to lack of proviral copies. (b) Representative flow cytometry data for Figure 2b demonstrating that GFP expression was restricted by the presence of both direct repeats (histogram overlays, first left). Complementary integration data depicting proviral copies in ESCs (second left). RNA was extracted from the experiments shown in Figure 2b and used for qRT-PCR on GFP mRNA to verify that epigenetic repression was at the transcriptional level consistent with TRIM28 epigenetic silencing. Data were normalized to expression of Cox6a1 mRNA (first right). GFP positive cells in 3T3 cells transduced with the same constructs as for ESCs for this figure (second right).

**Supplementary Figure 3.**
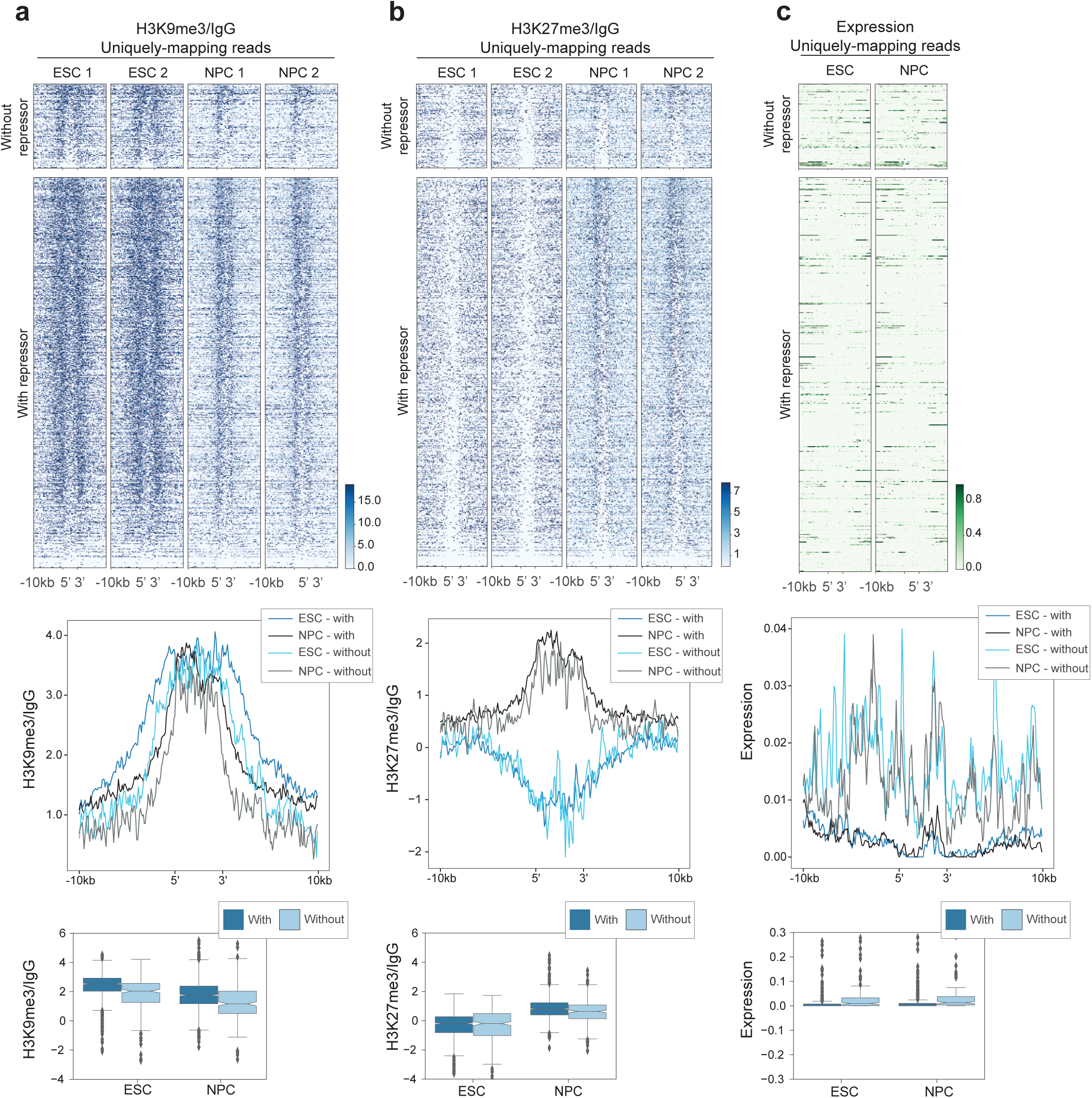
(a) H3K9me3 CUT&RUN signal normalised to IgG for uniquely-mapping reads across 838 IAPEz elements and surrounding 10Kb sequences to either end, grouped as with or without the repressor and sorted by decreasing intensity shown as a heatmap for two replicates (top). Profile plots depict the trimmed mean across all elements in the indicated category, across 100bp bins relative to the start and end coordinates of the element, where the body of IAPEzs is depicted in 50 bins (middle); boxplots show mean signal across the element and flanking regions for each IAPEz. Mann-Whitney U test, FDR corrected p value for ESC p=1.4e-18; NPC p=1.0e-15. (b) H3K27me3 CUT&RUN signal normalised to IgG using uniquely-mapping reads only, for two replicates as in (a), where elements in the heatmap are sorted by decreasing H3K9me3 signal. Mann-Whitney U test, FDR corrected p value for ESC p=0.84; NPC p=2.9e-05. (c) Mean RNA-Seq signal of uniquely-mapping reads across 3 replicates and normalised to number of mapped reads. Rows in the heatmap are sorted as in (a). Boxplots show mean signal across the element and 1000bp flanking regions for each IAPEz. Mann-Whitney U test, FDR corrected p value for ESC p=0.07; NPC p=0.08.

**Supplementary Figure 4.**
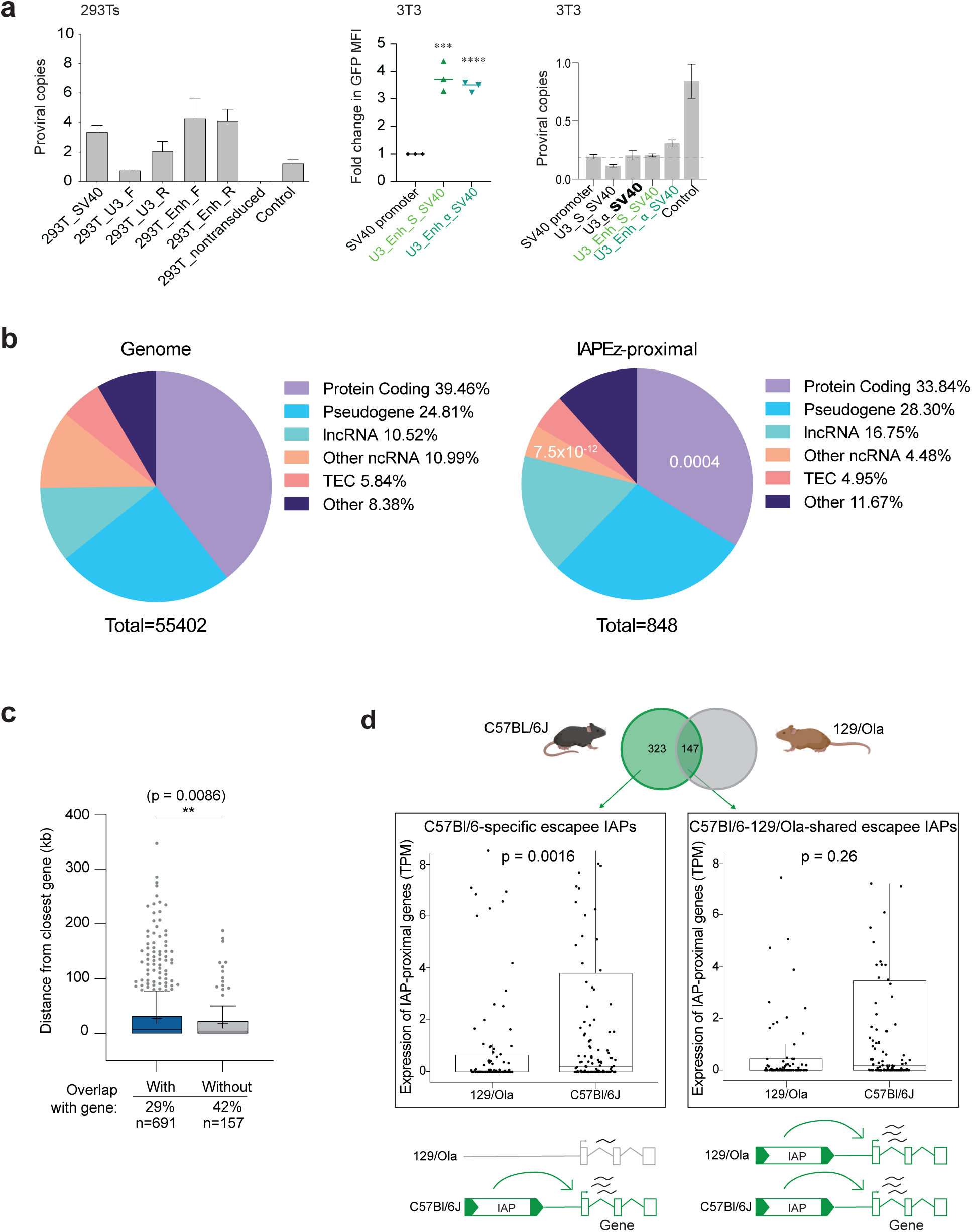
(a) Complementary data from Figure 4a to measure relative proviral integration in 293T cells and verify that the U3 enhancer acts as a *bona fide* enhancer rather than through differences in integration (left). GFP mean fluorescent intensity (MFI) and proviral copy integration for the same experiment as in Figure 4a carried out here in 3T3 cells (right). (b) Pie charts depicting the relative proportion of the listed gene biotypes (according to Ensembl) in the whole genome (left) compared to in a list of IAPEz-proximal genes (right) where hypergeometric tests were performed with P-values shown where difference was significant. (c) Boxplot of distances between IAPEz elements with or without a repressor sequence and their closest gene in kb, P-values calculated using Welch’s t-test, percentage of elements overlapping a gene as well as total number in each category is shown below. (d) Boxplots depicting TPMs of 323 genes closest to IAPEzs lacking a repressor sequence (escapee IAPs) which are present only in the C57Bl/6J strain as in Figure 4e and for 147 genes closest to escapee IAPEzs which are present in both the C57Bl/6J and 129/Ola strain where expression values are for 129/Ola NPCs or C57Bl/6 Cortical Neurons from (Bonev *et al*., 2017). P-values were calculated using a paired Wilcoxon test.

**Supplementary Figure 5.**
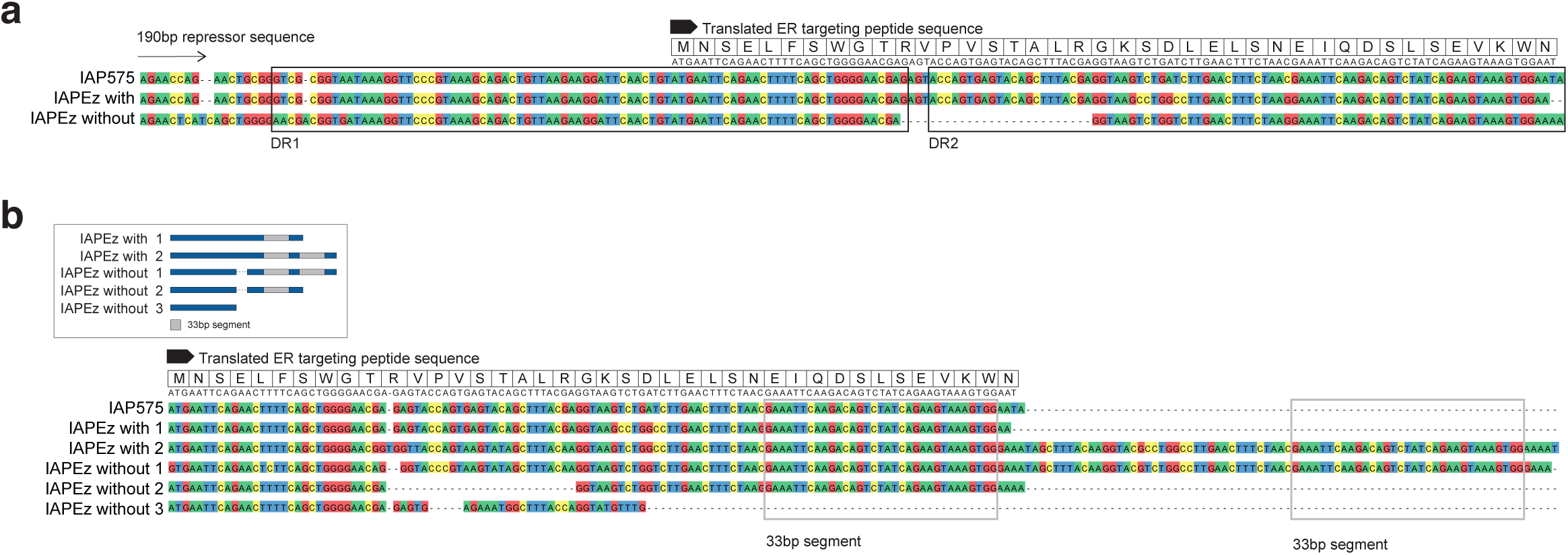
(a) Multiple sequence alignment of consensus IAPEz classified as with or without repressor obtained from manual annotation of the 190bp repressor sequence across IAPEz in the reference mouse genome and the canonical IAP575 repressor. The translated ER targeting peptide is shown above and both direct repeats are annotated in black. (b) Multiple sequence alignment (bottom) of consensus sequences of IAPEz subcategorized as depicted in box (top) and the canonical IAP575 sequence of the ER targeting sequence contained within the longer 190bp repressor sequence. Outlined in a grey box is a 33bp segment in direct repeat 2.

**Supplementary Information 1**

List of locations of IAPEz elements used in this study , their closest gene, presence or absence of repressor sequence and whether they are C57Bl/6 specific or shared between C57Bl/6 and 129/Ola strains.

## STAR METHODS

### RESOURCE AVAILABILITY

#### Lead contact

Further information and requests for resources and reagents should be directed to the Lead Contact, Helen Rowe.

#### Materials availability

Plasmids generated in this study are available on request by contacting the lead author.

#### Data and code availability

RNAseq and CUT&RUN sequencing data have been deposited at GEO and are publicly available as of the date of publication with accession numbers to be added.

This paper also analyzes existing, publicly available data. These accession numbers for the datasets are: GSE96107 (Bonev et al., 2017); GSE94323 (De Iaco et al., 2017).

Original western blot images have been deposited at Mendeley and are publicly available as of the date of publication. DOI to be added.

All original code has been deposited at GitHub and is publicly available as of the date of publication; DOIs to be added.

Any additional information required to reanalyze the data reported in this paper is available from the lead contact upon request.

## EXPERIMENTAL MODEL AND SUBJECT DETAILS

### Cell culture and reagents

ES3 mouse embryonic stem cells were derived from C57Bl/6 mouse embryos (male) and were a gift from Didier Trono (Rowe et al., 2010). 46C ESCs (Ying et al., 2003) are a Sox1-GFP reporter cell line generated by gene targeting of E14Tg2a.IV ES cells which are derived from a 129/Ola mouse embryo (male) and were a gift from Prof. Austin Smith (University of Cambridge, UK). Mouse embryonic stem cells were cultured in N2B27 media: DMEM/F12 (Gibco, Thermo Fisher Scientific ), Neurobasal (Gibco, Thermo Fisher), N2 (Gibco, Thermo Fisher), B27 (Gibco, Thermo Fisher), 0.1mM 2-Mercaptoethanol (Life Technologies) and supplemented with 0.08% BSA and 100 U/ml penicillin/streptomycin (Pen-Strep, Life Technologies), under 2i/LIF culture conditions: 1,000units/ml Leukaemia inhibitory factor (LIF, Chemicon), 1μM PD0325901 (Merck, Sigma Aldrich) and 3μM CHIR99021 (Merck, Sigma Aldrich). Cells were grown at 5% CO2 at 37°C and split 1:4 every 2 days with Accutase. 3T3 and HEK293T cells were cultured in Dulbecco’s Modified Eagle’s Medium (DMEM; Gibco, Thermo Fisher Scientific) with 10% FBS and 100 U/ml penicillin/streptomycin, grown at 5% CO2 and split every 1:5 every 2 days with trypsin.

## METHOD DETAILS

### Plasmids and lentiviral vectors

Locus-specific cloning of the IAP sequences described in the text into a lentiviral GFP vector in place of the hPGK promoter was carried out as described previously (Rowe *et al*., 2013a) and products were verified by sequencing. Here, the IAP LTR from the IAP575 locus served as the promoter driving GFP expression itself rather than an internal promoter as previously used (Rowe *et al*., 2013a). Flow cytometry was used to measure GFP expression for the reporter assays. For RNAi *Trim28* pLKO.1 hairpins were purchased from Dharmacon and the empty pLKO.1 vector used as control. Cells were selected with puromycin overnight (or until control cells had all died) before collection and analysis. Lentiviral vectors were produced by Fugene6 co-transfection of HEK293T cells with 1.5μg of plasmid, 1μg p8.91 and 1μg pMDG2 encoding VSV-G in a 10cm plate. Ultracentrifugation of the supernatant (20,000g for 2h at 4°C) was carried out 2 days post transfection.

### Intracellular POU5F1 staining/SSEA1 staining

ESCs were fixed and surface stained with anti-SSEA1-PerCP and then permeabilized using the eBioscience intracellular staining buffer kit (eBioscience 88-8824-00) and subjected to intracellular staining with anti-POU5F1-PE, or isotype control antibody and analysed by flow cytometry. See Supplementary Table 1 for antibody information.

### RNA extraction and quantification

Total RNA was extracted using an RNeasy micro kit (Qiagen) and treated with DNase (Ambrio, AM1907). cDNA was synthesised from 500ng of RNA with SuperScript II Reverse Transciptase (ThermoFisher Scientific) using random primers. RT-qPCR was carried out using SYBR green Fast PCR mastermix (Life Technologies) on an ABI 7500 Real Time PCR System (Applied Biosystems). CT values were normalised to *Cox6a1* and fold change was calculated using the -DDCt method. See Supplementary Table 2 for primer sequences.

### Western Blotting

Cells were washed in PBS and lysed in cold RIPA buffer (150 mM NaCl; 1% Triton X-100; 0.5% sodium deoxycholate; 0.1% SDS and 50 mM Tris, pH 8.0, and protease inhibitor cocktail (cOmplete™, Mini, EDTA-free, Roche)), lysates were quantified for normalisation (BCA Protein Assay kit, Millipore) and loaded on 10% denaturing SDS-polyacrylamide gels. Wet transfers were carried out onto PVDF membranes, blocked in 5% milk in TBS-T (TBS, 0.1% Tween-20 (Sigma)) and incubated with antibodies. Membranes were visualised using Amersham ECL kits. Antibodies used were: anti-PCNA, anti-POU5F1, anti-TRIM28, anti-Nanog. See Supplementary Table 1 for antibody information.

### NPC differentiation

46C ESCs were cultured in 2i/LIF conditions as described above and then cultured for two passages without LIF; NPCs were generated from these cells as described in (Gouti et al., 2014), with some modifications: ESCs were plated on laminin-coated 6 well plates at a density of 65000 cells per well in N2B27 media (as above but with N2 Supplement-B from StemCell Technologies) supplemented with bFGF (10ng/uL bFGF (R&D)) and 1μg/mL laminin. Cells were cultured for 5 days with daily media changes (day 1-2 with bFGF, day 3-5 without bFGF) at 7% CO2 and at day 5, cells were collected and analysed by flow cytometry (ACEA Novocyte 3000) to measure GFP expression and used for downstream analyses.

### CUT&RUN

CUT&RUN was carried out according to the EpiCypher® CUTANA CUT&RUN Protocol (v1.6) (https://www.epicypher.com/resources/protocols/cutana-cut-and-run-protocol/). 100,000 ESCs or day 5 NPCs were collected per sample/antibody and washed twice with EpiCypher Wash buffer (20mM HEPES pH 7.5, 150mM NaCl, 0.5mM spermidine) plus protease inhibitors (cOmplete™, Mini, EDTAfree, Roche) before attachment to activated Concanavalin A coated magnetic beads. Beads and cells were resuspended in Antibody Buffer (20mM HEPES pH 7.5, 150mM NaCl, 0.5mM Spermidine, 1x protease inhibitors, 0.01% w/v digitonin, 2mM EDTA) with antibodies (1ug of anti-H3K9me3, anti-H3K27me3 or IgG control; Supplementary Table 1) and incubated overnight at 4°C with gentle rocking. The next day beads were washed twice in cold Digitonin Buffer (20 mM HEPES pH 7.5, 150mM NaCl, 0.5mM Spermidine, 1x Roche complete protease inhibitors, 0.01% digitonin) and then incubated with Digitonin Buffer plus 2.5μL pAG-MNase (CUTANA™, Epicypher) before addition of 2mM CaCl_2_ to activate cleavage at 4°C for 2h. The reaction was quenched by addition of 33μl Stop Buffer (340mM NaCl, 20mM EDTA, 4mM EGTA, 50μg/ml glycogen, 50μg/ml RNase A), vortexted and incubated at 37°C for 10minutes to enable the release of DNA fragments. Sample was cleaned using a magnetic rack and the supernatant containing DNA was purified with a MinElute PCR Purification Kit (Qiagen). Libraries were prepared according to manufacturer’s instructions using the following kits: NEBNext® Ultra™ II DNA Library Prep Kit for Illumina® and NEBNext® Multiplex Oligos for Illumina® (New England BioLabs) and pooled in equimolar quantities. Sequencing was carried out on a Novaseq6000 with 150bp PE reads.

### CUT&RUN data analyses

Reads were trimmed and adapters removed using TrimGalore v0.4.1 (Krueger; https://www.bioinformatics.babraham.ac.uk/projects/trim_galore/) quality was checked using FastQC v.0.11.8 (Anders). Alignment to the GRCm38 mouse genome was performed using STAR (Dobin et al., 2013) with parameters adjusted to generate one random location for multimapping reads *[--outFilterMultimapNmax 5000 --outSAMmultNmax 1 -- outFilterMismatchNmax 999*]. The genomecov tool from bedtools was used to generate BedGraph files of genome coverage, scaled by library size, and converted to BigWig files. The pybigwig library in python3 was used to retrieve coverage over 100bp windows. CUT&RUN signal was normalised to IgG signal, this data was represented at heatmaps, profile plots and boxplots using the matplotlib and seaborn libraries.

Alignments of uniquely mapping reads were also performed with STAR with the *-- outFilterMultimapNmax 1* option and processed as above.

### ChIP-Seq data analysis

Data from (De Iaco *et al*., 2017) (GSE94323), was downloaded using the SRA toolkit. STAR alignments of reads were performed as above, allowing for one random location for multimapping reads. Alignments were processed similarly as CUT&RUN data to generate genomewide coverage in bigwig format for IP and input samples. IP signal was normalised to input in python3 using the pybigwig library and depicted was profile plots or heatmaps as indicated.

### RNA sequencing

RNA was extracted as described above and RNA quality and quantity was assessed using a Spectrophotometer UV5 (Mettler Toledo). Preparation of mRNA libraries was carried out by Novogene Co. Ltd and sequenced on the Illumina NovaSeq6000 with 150bp PE reads. Data was demultiplexed and fastq files generated using the bcl2fastq software from Illumina. TrimGalore v0.4.1 (https://www.bioinformatics.babraham.ac.uk/projects/trim_galore/) was used for read trimming and adapter removal and FastQC v.0.11.8 (Anders et al., 2010) was used for quality checking. Reads were aligned to the GRCm38 mouse genome using STAR *[-- outFilterMultimapNmax 5000 --outSAMmultNmax 1 --outFilterMismatchNmax 999*]. Read counts per gene were obtained with HTSeq-Count (Anders et al., 2015) and differential expression analysis was performed with DESeq2 (Love et al., 2014) under R v4.1.1 was used to call differential expression analysis for genes, and the Approximate Posterior Estimation method (Zhu et al., 2019) was used to shrink the logarithmic fold change. TPM values were calculated in R. For depiction of expression signal, genome coverage scaled by library sized was calculated with the genomecov tool of bedtools and converted to bigwig to be processed in python3 as above. Data from (Bonev *et al*., 2017) (GSE96107) were downloaded using the SRA toolkit and the reads mapped with STAR and TPMs were calculated as above.

### Functional analysis of genes

bedtools closest tool (v2.27.1) was used to call the closest gene and distance to each IAP. GO analysis was performed on the list of closest genes using the Bioconductor clusterprofiler tool (Wu et al., 2021) in R. Genes were classified according to their transcript biotype where this information was collected using the Bioconductor biomaRt tool (Durinck et al., 2009) in R.

### Sequence analyses

The intersect and closest tools from bedtools were used to obtain full-length IAPEz elements which complied with the following: contained two IAPLTR1 or IAPLTR1a, in the same orientation and were separated by less than 20kb. We then verified that elements which fulfilled the previous also overlapped a ‘IAPEz’ internal sequence. The sequences corresponding to these elements were extracted with the getfasta tool from bedtools. Matches to the 190bp repressor sequence of IAP575 were calculated with the water tool from EMBOSS *[-gapopen 10 -gapextend 0.5*] and the %identity was used to classify elements based on their repressor sequences. Multiple sequence alignments were generated with muscle (Edgar, 2004) and visualized using Jalview (Waterhouse et al., 2009). Alignments were manually curated to subcategorise repressor types and consensus sequences were generated with HMMER (Eddy, 2011) using the hmmbuild and hmmemit tools.

### Quantification and statistical analysis

Data shown in this study are shown with error bars representing standard deviation. Where shown statistitcal significance was assessed with two tailed, paired Student’s t tests or as described in the figure legends using GraphPad Prism or R v4.1.1. Biological replicates are denoted in the figure legends. For flow cytomentry 10,000 events were recorded. P-values of <0.05 were considered significant (****P < 0.0001, ***P < 0.001, **P < 0.01 and *P < 0.05) and P-values are shown in the figure or legends.

## TABLES

**Supplementary Table 1:** List of antibodies

| Protein | Protocol | Species | Cat. No. | Manufacturer |
| --- | --- | --- | --- | --- |
| OCT4 | FACS | Rat | 12-5841 | eBioscience |
| SSEA1 | FACS | Mouse | eBioMC-480 | eBioMC |
| PCNA | WB | Mouse | NA03 | Calbiochem |
| OCT4 | WB | Mouse | sc-5279 | Santa Cruz Biotech |
| TRIM28 | WB | Rabbit | Ab10483 | Abcam |
| NANOG | WB | Rabbit | Ab80892 | Abcam |
| H3K9me3 | C+R | Rabbit | Ab8898 | Abcam |
| H3K27me3 | C+R | Rabbit | Ab195477 | Abcam |
| IgG control | C+R | Rabbit | 12-370 | Millipore |
| IgG control | FACS | Rat | 12-4321 | eBioscience |

**Supplementary Table 2:**
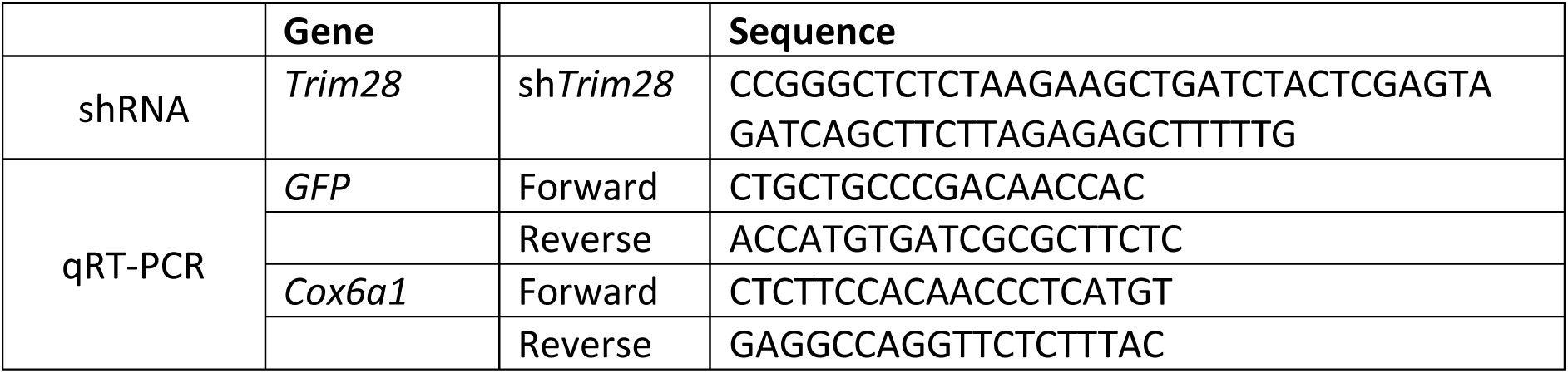
shRNA and primer sequences

